# Restriction Endonuclease-based Modification-Dependent Enrichment (REMoDE) of DNA for Metagenomic Sequencing

**DOI:** 10.1101/2022.09.30.510419

**Authors:** Syed Usman Enam, Joshua L. Cherry, Susan R. Leonard, Ivan N. Zheludev, David J. Lipman, Andrew Z. Fire

## Abstract

Metagenomic sequencing is a swift and powerful tool to ascertain the presence of an organism of interest in a sample. However, sequencing coverage of the organism of interest can be insufficient due to an inundation of reads from irrelevant organisms in the sample. Here, we report a nuclease-based approach to rapidly enrich for DNA from certain organisms, including enterobacteria, based on their differential endogenous modification patterns. We exploit the ability of taxon-specific methylated motifs to resist the action of cognate methylation-sensitive restriction endonucleases that thereby digest unwanted, unmethylated DNA. Subsequently, we use a distributive exonuclease or electrophoretic separation to deplete or exclude the digested fragments, thus, enriching for undigested DNA from the organism of interest. As a proof-of-concept, we apply this method to enrich for the enterobacteria *Escherichia coli* and *Salmonella enterica* by 11- to 142-fold from mock metagenomic samples and validate this approach as a versatile means to enrich for genomes of interest in metagenomic samples.

**Importance:** Pathogens that contaminate the food supply or spread through other means can cause outbreaks that bring devastating repercussions to the health of a populace. Investigations to trace the source of these outbreaks are initiated rapidly but can be drawn out due to the labored methods of pathogen isolation. Metagenomic sequencing can alleviate this hurdle but is often insufficiently sensitive. The approach and implementations detailed here provide a rapid means to enrich for many pathogens involved in foodborne outbreaks, thereby improving the utility of metagenomic sequencing as a tool in outbreak investigations. Additionally, this approach provides a means to broadly enrich for otherwise minute levels of modified DNA which may escape unnoticed in metagenomic samples.

## Introduction

Foodborne pathogen outbreaks can be a major public health and agro-economic burden (1, 2). According to the World Health Organization, one in ten people are victim to foodborne illnesses every year (https://www.who.int/news-room/fact-sheets/detail/food-safety). When such outbreaks occur, food and agricultural safety organizations are tasked with determining the responsible contaminated food, the pathogen causing the illness and the source of this pathogen so that required measures can be taken to remove implicated food products from commerce and perform remediation steps to prevent further illnesses (1–5).

Specific strains of enterobacteria are amongst the common pathogens linked to foodborne illnesses (e.g. *Salmonella enterica*, Shiga toxin-producing *Escherichia coli*) (1–7). In an outbreak setting, pathogen detection and identification are often achieved through serotype testing, DNA marker amplification, or targeted sequencing of genomic loci, but these methods sometimes provide insufficient information to trace the organism back to its source (1, 3–5, 7, 8). Thus, strain and sub-strain level information encoded by single nucleotide polymorphisms (SNPs) can have great value in tracking and matching a pathogen to its environmental source (1, 9). Whole genome sequencing (WGS), unlike targeted methods, provides this information and is used at various checkpoints between the farm and consumer to monitor and control contamination in produce (1–5, 10).

WGS of an outbreak pathogen is obtained through either a culture-dependent or a culture-independent approach, the first of which can add many days and substantial cost to an investigation depending on how simple it is to isolate the pathogen in question and the pathogen load in the sample (1, 3–5, 7). In a culture-independent approach, shotgun sequencing is performed on the sample (produce, food, soil, plant) potentially containing the pathogen and assembly of the pathogen’s DNA allows for rapid strain-level identification without the need for isolation. While shotgun WGS is recognized as a powerful tool for this application, it comes with the limitation that a sample needs to contain a sufficient load of the pathogen for high enough coverage of the genome to SNP map the genome. Often these samples, instead, contain irrelevant prokaryotic and eukaryotic DNA that far exceeds that of the relevant strain. This leads to low-coverage assemblies of the pathogen or increased cost due to excess sequencing of a sample (11–16).

Methods exist to deplete “host” or eukaryotic DNA and enrich for prokaryotic DNA. Some commercial kits selectively lyse eukaryotic (mostly mammalian) cells and degrade accessible DNA through enzymatic or chemical means before purifying DNA from the remaining cells (4, 14, 15). While these offer substantial depletion of eukaryotic DNA, they may fall short in several ways: 1) unable to digest (and therefore deplete) cells with robust cell walls such as fungi, 2) unable to enrich for prokaryotic DNA post-DNA extraction, 3) unable to preserve prokaryotic cell-free DNA in a sample and 4) unable to deplete irrelevant prokaryotic DNA.

Other methods deplete eukaryotic DNA post-DNA extraction by binding and sequestering this DNA due to the differential presence of methylation patterns between prokaryotes and eukaryotes (11–13, 15, 16). One commercial kit takes advantage of the increased presence of CpG (C5 position of cytosine) methylation in eukaryotes and uses an engineered methyl-CpG binding domain conjugated to an antibody to remove CpG methylated DNA (16). However, studies frequently report weak enrichment through this method likely due to the presence of large stretches of eukaryotic DNA that are not methylated (14, 15). Additionally, many eukaryotes, such as *Caenorhabditis elegans*, exhibit predominantly unmodified DNA (17, 18). A different protocol makes use of methylation-sensitive restriction enzyme HpaII in non-catalytic conditions to bind and enrich for non-CpG methylated (and therefore prokaryotic) DNA (12). The same group also applied this paradigm to a different restriction enzyme, DpnI which selectively targets methylated 5’-GATC-3’ motifs (N6 position of adenine) (11). These motifs are methylated by Dam, a type-II methyltransferase widespread in *Gammaproteobacteria* (of which *E. coli, S. enterica* and *Vibrio cholerae* are members) and not found in eukaryotes (19–24). While offering substantial enrichment, these protocols are time- and cost-prohibitive as they involve using 1:1 stoichiometric amounts of enzyme to the to-be-enriched substrate DNA, modification of the enzyme by biotinylation and a final dialysis step (11, 12).

Here we describe and implement REMoDE: Restriction Endonuclease-based Modification-Dependent Enrichment of DNA, an approach that rapidly and cost-effectively enriches for DNA from *E. coli* and *S. enterica* in metagenomic samples. We, too, rely on the presence of Dam and Dcm methyltransferases in *E. coli* and *S. enterica* and the near complete methylation of all instances of their target motifs in *E. coli* and *S. enterica* (23–25). These methyltransferases methylate 5’-G<u>**A**</u>TC-3’ and 5’-C<u>**C**</u>WGG-3’ respectively (methylated base underlined; W = A or T (26)) (23, 25, 27). We employ the highly specific action of methylation-sensitive restriction enzymes MboI (5’-GATC-3’), PspGI and EcoRII (both 5’-CCWGG-3’) that cleave only unmethylated instances of the motif (28). When applied to a mixed population of DNA that is unmethylated or methylated at these motifs, we observe a segregation of DNA into either short or long, genomic-length fragments respectively. Finally, we select for the longer fragments of DNA either using electrophoretic separation or by taking advantage of the highly distributive nature of the T5 exonuclease (29). When applied to a reaction with different distributions of long and short DNA, electrophoretic separation provides a clean size separation, albeit requiring an additional gel isolation step, while the T5 exonuclease reaction is a cost-effective approach (~$0.03 additional-cost/sample) that can be adjusted to rapidly deplete short DNA in a same tube reaction. We observe a 11- to 142-fold enrichment of *E. coli* and *S. enterica* DNA in a metagenomic sample relative to an untreated version of the same sample. This method can be extended to other Dam and Dcm methylated organisms and may even be extrapolated to organisms with other methylation patterns as pointed out in the discussion.

**Fig 1:**
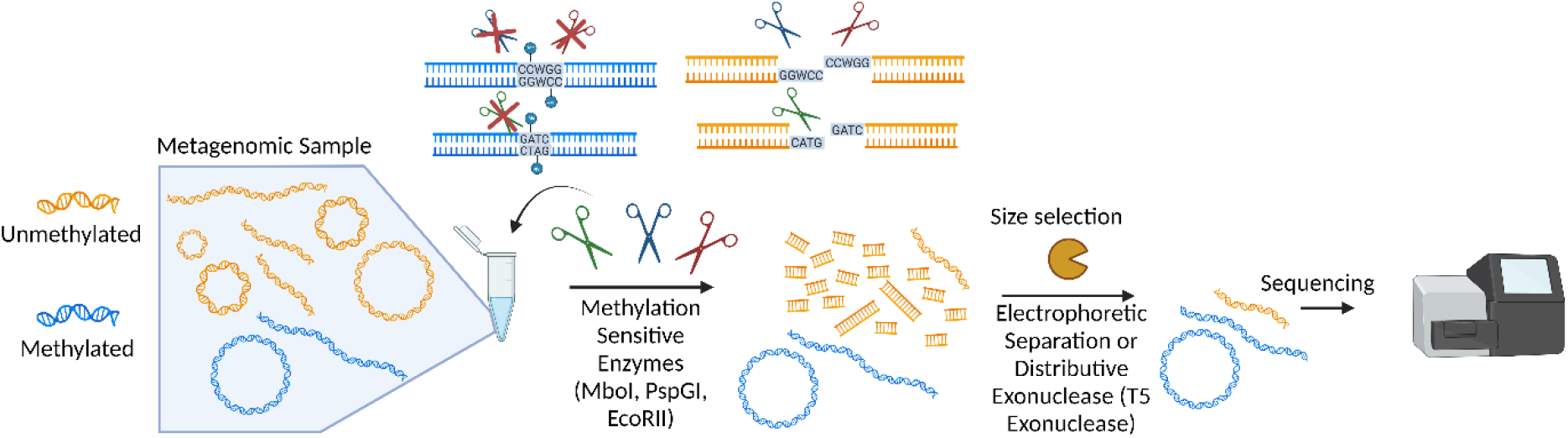
Schematic of the pipeline for endonuclease and exonuclease-based enrichment of methylated DNA. A metagenomic sample containing DNA that is and is not Dam and Dcm methylated is treated with methylation sensitive enzymes. The unmethylated DNA is digested to short fragments while the methylated DNA remains long and intact. Size selection for longer fragments is performed with either electrophoretic separation or a distributive exonuclease (which preferentially degrades short fragments). The enriched sample is then sequenced.

## Results

Figure 1 provides an overview of the restriction-enzyme-based scheme that we have used to enrich for DNAs methylated at defined sites.

As a proof of principle, we elected to test this approach with DNA from organisms readily available in the laboratory and that we knew were Dam and Dcm methylated (*E. coli*) or unmethylated (*C. elegans*).

Genomic DNA from TOP10 *E. coli* and N2 *C. elegans* was prepared and treated with restriction endonucleases MboI, PspGI and EcoRII. The DNA was found to be either resistant or susceptible to the action of these endonucleases, respectively (Fig 2A). A 1:3 mixture (by mass) of genomic DNA from *E. coli* and *C. elegans* was prepared as a stand-in for a metagenomic sample. After treatment with the endonucleases, the sample was treated with the distributive T5 exonuclease for 2, 5, 10 or 20 minutes. When treated for five minutes, or beyond, shorter fragments were substantially depleted from the sample, while longer fragments were retained (Fig 2B). The preferential retention of longer fragments was as expected given the timing of the reaction and rates of terminus-degrading exonuclease activity (29). When sequenced on an Illumina MiSeq, a progressive enrichment of the proportion of reads that mapped to *E. coli* was observed (Fig 2C). Relative enrichment values were calculated as the ratio of the number of *E. coli* reads to *C. elegans* reads in a treated sample divided by the ratio in an untreated sample. With 20 minutes of exonuclease treatment, a 27.5 fold enrichment was observed (Fig 2C). Longer T5 treatment did not result in greater enrichment of *E. coli* reads in this mixture (different experiment; data not shown). To determine where the remaining proportion of *C. elegans* reads were originating from, each *C. elegans* read was assigned the theoretical length of the restriction fragment it came from in an *in-silico* digestion of the *C. elegans* genome. The cumulative distribution of these lengths was plotted for each T5 exonuclease time point (Fig 2D) and many *C. elegans* reads, as expected, originated from regions greater than 10kb.

**Fig 2:**
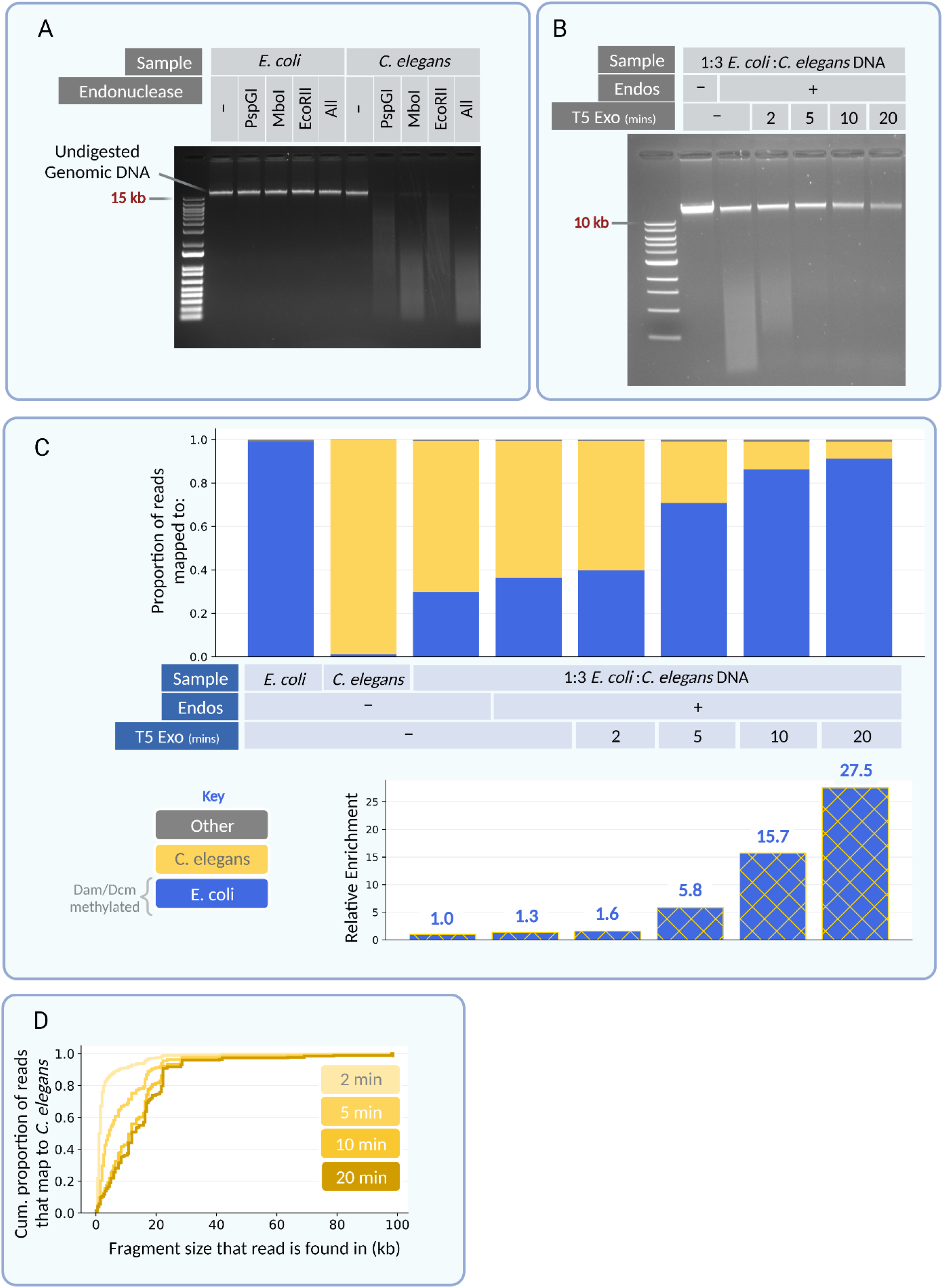
Methylation sensitive endonucleases and T5 exonuclease enrich for *E. coli* DNA in an *E. coli* and *C. elegans* DNA mixture. (A) Gel showing the susceptibility of either E. coli DNA or C. elegans DNA to PspGI, MboI and EcoRII separately and all together. The genomic high molecular weight C. elegans band disappears when the endonuclease is applied. (B) Gel showing timepoints of T5 exonuclease treatment when applied to a 1:3 mixture of E. coli to C. elegans DNA treated with the corresponding endonucleases. Notice the disappearance of the low molecular weight smear (C. elegans DNA) with longer T5 exonuclease incubation. (C) Paired end sequencing data from untreated and treated samples. In blue are the proportion of reads in that sample that map to the E. coli genome and in yellow are the proportion that map to the C. elegans genome. Any reads that do not map to either or have chimeric paired reads are colored grey. The C. elegans only sample contains a certain amount of E. coli DNA likely due to the fact that the worms are fed E. coli. Shown below is the relative enrichment of E. coli DNA calculated as the ratio of the number of E. coli reads to C. elegans reads divided by the ratio in the untreated control. (D) For each T5 exonuclease time point, all C. elegans reads that remained were mapped to the length of the theoretical fragment size that they would be found in an in silico digestion of the C. elegans genome. A cumulative density plot of these fragments is shown to ascertain whether remaining C. elegans reads originate from long fragments or short fragments.

To assay a more complex but standardized sample, we performed this treatment on ZymoBIOMICS microbial community standard high molecular weight DNA (MCS-HMW). This is a mixture of genomic DNA from one yeast and seven bacteria – of which two (*E. coli* and *S. enterica*) are Dam and Dcm methylated. Upon PspGI, MboI and EcoRII endonuclease and T5 exonuclease treatment, an average of a 10.8-fold relative enrichment of *E. coli and S. enterica* DNA was observed (Fig 3A). DNA from these two species composes 28% of the untreated MCS-HMW mixture according to the manufacturer (Zymo Research). However, we found that roughly 35 to 40% of reads from an untreated sample map to the *E. coli* and *S. enterica* genomes. This is likely due to our transposition-based sequencing library construction methods which have a bias against GC-rich genomes (30). Of note, also, is that five minutes of T5 exonuclease treatment resulted in greater enrichment of *E. coli* and *S. enterica* DNA than 20 minutes of T5 exonuclease treatment (data not shown) unlike what was seen with the mixture of *C. elegans* and *E. coli* DNA. This may have been due to the different ways the genomic DNA was prepared as well as the number of free ends available in a given exonuclease reaction. Additionally, it was observed that the enrichment worked just as well without EcoRII, which thus may be omitted (Fig S1). However, to maintain experimental consistency, EcoRII was used for all following experiments.

**Fig 3:**
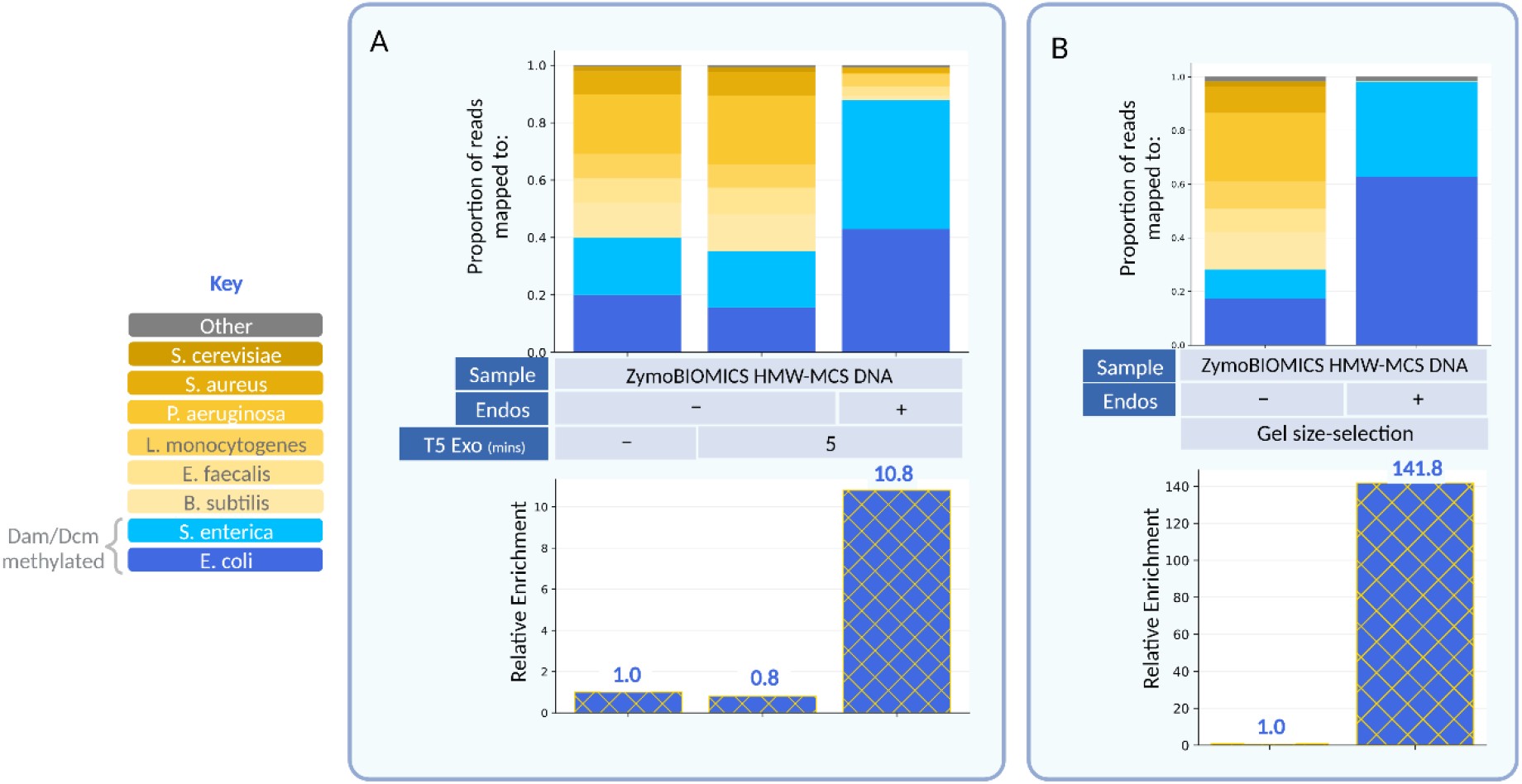
Methylation sensitive endonucleases and various size-selection approaches enrich for *E. coli* and *S. enterica* DNA in the ZymoBIOMICS MCS-HMW DNA. (A) Paired end sequencing data from untreated, endonuclease-only treated and endonuclease as well as T5 exonuclease treated DNA. In blue are reads that map to genomes that are Dam and Dcm methylated. In yellow are reads that map to genomes that are not Dam and Dcm methylated. Relative enrichment of E. coli and S. enterica DNA shown below. (B) Paired end sequencing data from untreated and endonuclease treated DNA size-selected through gel electrophoresis (Fig S2). Relative enrichments are shown below. Means of two biological duplicates were plotted for (A) and (B).

The addition of the T5 exonuclease acts to select for long fragments of DNA rapidly (5 to 20 mins). This approach has the advantage of a low cost (~$0.03/sample) and can be performed in the same tube as the endonuclease treatment. We were curious how this might compare to the gold standard of electrophoretic size selection (agarose gel electrophoresis). Endonuclease untreated and treated MCS-HMW DNA samples were resolved on a gel alongside each other (Fig S2). Due to the size exclusion limit of a 1% gel, all fragments greater than 15 kb (highest band of the ladder) comigrate as a single band (31). In the treated sample, a smear throughout the lane is observed, however high molecular weight DNA originating from *E. coli* and *S. enterica* is found near the 15 kb mark. Both the untreated and the treated high molecular weight bands were extracted from the gel and sequenced. Virtually all of the reads from the treated sample mapped to *E. coli* and *S. enterica*, providing a substantial relative enrichment of 141.8-fold (Fig 3B). Electrophoretic size selection therefore proves to be an effective way of separating digested fragments from undigested fragments. However, it comes at the cost of time and money over a T5 exonuclease size selection.

Next, to test the dynamic range of this enrichment, we titrated down the input MCS-HMW DNA amount from the initial value of 75ng to 37.5ng (1/2), 7.5ng (1/10) and 0.75ng (1/100). In all cases, we observed on average a greater than 20-fold relative enrichment of *E. coli* and *S. enterica* DNA (Fig 4A).

**Fig 4:**
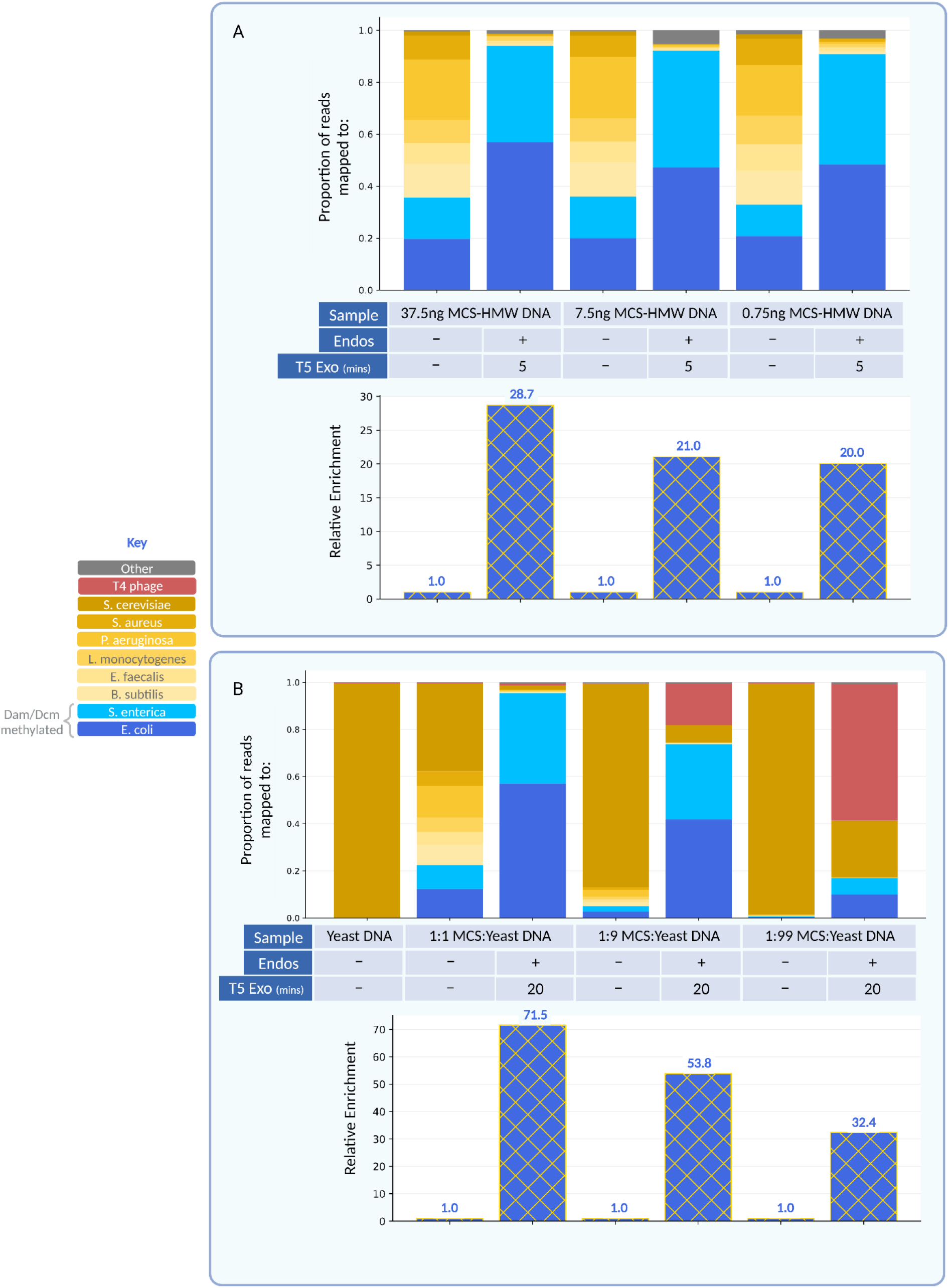
Dynamic range of enrichment on various amounts and ratios of methylated DNA. (A) Paired end sequencing data from either half, one-tenth or one-hundredth the amount of ZymoBIOMICS MCS-HMW DNA used in the standard protocol following otherwise the same enzyme concentrations. In blue are reads that map to genomes that are Dam and Dcm methylated. In yellow are reads that map to genomes that are not Dam and Dcm methylated. Relative enrichment is shown below. (B) Paired end sequencing data from ZymoBIOMICS MCS-HMW DNA mixed with S. cerevisiae in 1:1, 1:9 and 1:99 ratios with total amount remaining 75ng. Means of two biological duplicates were plotted for (B). In red are reads that map to T4 phage. T4 phage DNA sequences represent a population of DNA fortuitously included with the yeast DNA material used in these assays. Notably, this DNA is enriched in parallel to the modified bacterial DNAs, a behavior that is both of interest and expected as a consequence of the known modification of T4 DNA. Thus, this population serves as a fortuitous positive control on the enrichment observed.

While the protocol continued to be effective for vanishingly small amounts of input DNA, we questioned how the enrichment varied when the ratio of Dam/Dcm methylated DNA to unmethylated DNA was altered. To test this, the ZymoBIOMICS MCS-HMW DNA was mixed with *Saccharomyces cerevisiae* genomic DNA in a 1:1, 1:9 and 1:99 ratio. We observed a robust enrichment of *E. coli* and *S. enterica* DNA at all ratios, with average enrichment ranging from 32.4- to 71.5-fold (Fig 4B). To note, is the surprising enrichment of a class of reads that did not seem to map to any of the genomes present in the ZymoBIOMICS MCS-HMW standard or *S. cerevisiae*. This class of reads was reproduced upon replicate experiments. These reads were assembled into contigs using SPADES (32). The largest, most prevalent contig from these unmapped reads mapped to *E. coli bacteriophage T4* using BLAST (shown now as red in Fig 4B). Upon closer inspection, we realized some phage T4 DNA was unintentionally included with the *S. cerevisiae* DNA. Enrichment of T4 DNA is expected because it contains hydroxymethyl cytosine, usually glucosylated, in place of cytosine (33–35). This allowed the T4 DNA to resist cleavage by PspGI, MboI and EcoRII and was therefore selected for during T5 exonuclease treatment, serving as a fortuitous positive control. This happenstance points towards the ability to use this tool as way to discover genomes in metagenomic samples that are substantially modified or contain non-canonical bases (see discussion).

Finally, we asked whether a parallel protocol could be used for selective enrichment of unmodified DNA. *DpnI* is a restriction endonuclease that selectively cleaves at Dam sites that are *methylated* (unlike MboI which cleaves at Dam sites that are unmethylated) (24). Accordingly, DpnI can be used to deplete *E. coli* and *S. enterica* DNA in a metagenomic sample. When DpnI was applied to the ZymoBIOMICS MCS-HMW DNA, a 7.6-fold relative enrichment of non-Dam methylated DNA was observed as compared to the untreated control (Fig 5).

**Fig 5:**
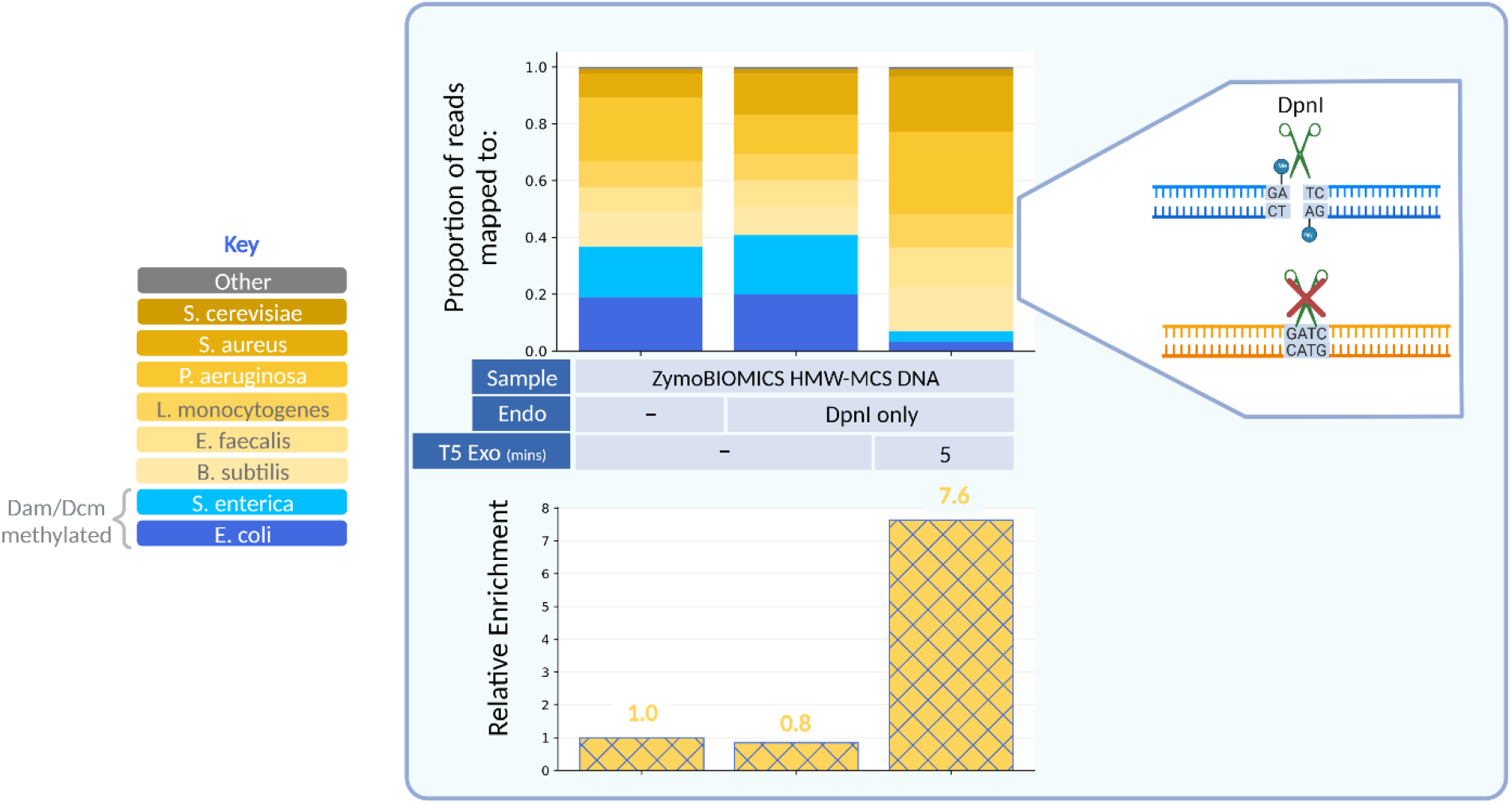
DpnI and T5 exonuclease treatment enriches for DNA that is not Dam methylated. Paired end sequencing data for ZymoBIOMICS MCS-HMW DNA treated with DpnI which only cuts at methylated Dam sites. Relative enrichments are shown below.

## Discussion

In this work we have described and implemented a novel approach, REMoDE, to enrich metagenomic samples for DNA from organisms of interest based on their specific patterns of DNA modification. While differential methylation has been used to obtain enriched sequence datasets in the past, the technical approaches have involved binding and release steps with high complexity in terms of reagents and protocols (11, 12). Applying restriction enzyme cleavage followed by exonuclease- or gel-based size selection, we obtained remarkable enrichments with only limited manipulation.

The approach provided herein specifically implements methods to selectively enrich DNA of organisms that contain Dam and Dcm systems. These methyltransferases are found in many members of the *Gammaproteobacteria* phyla including *E. coli* and *S. enterica* (19). Many pathogenic food outbreaks have been caused by species from the *Gammaproteobacteria* phyla (2). Various food and agricultural safety applications require high sequencing coverage of the outbreak strain to confidently obtain identifying SNPs for an outbreak source (optimal coverages may be as high as 50x (36)). Such coverage allows potential matching of the agricultural source with contaminated foods, providing an opportunity to accurately restrict further outbreak from the source, while avoiding interference with supply chains uninvolved in an outbreak.

Sequencing approaches lend tremendous specificity and sensitivity to detection and characterization of potential pathogens. However, challenges in utilization of a sequencing approach can arise in that metagenomic samples from both environmental and clinical sources generally contain irrelevant prokaryotic and eukaryotic DNA that far exceeds that of the pertinent strain and therefore obtaining high coverage WGS can prove difficult. This approach proves especially useful in metagenomic analyses such as these. In our experiments, we observe enrichment of *E. coli* and *S. enterica* DNA ranging from 11-fold to 142-fold with a broad dynamic range for input DNA amount. Additionally, the method has been developed such that the enrichment can be performed in a single tube and completed within one-and-a-half hours.

Dam and Dcm systems extend to clinically relevant organisms beyond *E. coli* and *S. enterica* that would benefit from whole genome sequencing for tracing purposes (19). For example, *Vibrio cholerae* (causes cholera), *Yersinia pestis* (causes plague), *Legionella pneumophila* (causes Legionnaire’s disease), and *Klebsiella pneumoniae* (cause pneumonia) are either known or predicted to methylate their Dam sites (19).

Some eukaryotic viruses are also known to harbor methyltransferases. For example, the Melbournevirus of the giant virus family Marseilleviridae is also Dam methylated (37).

It is also worth noting that the principle of enrichment using restriction enzymes and an exonuclease need not only extend to Dam and Dcm methylated DNA. As shown, unmodified DNA can be enriched using DpnI and this paradigm can be applied more broadly by taking advantage of the vast catalogue of restriction enzymes. Indeed, among others there are two methylated motifs that are broadly prevalent in clinically relevant bacteria: 5’-RA<u>**A**</u>TTY-3’ and 5’-G<u>**A**</u>NTC-3’ (R = A or G; Y = C or T; N = any (26)) (19). Unmethylated 5’-RAATTY-3’ is endonuclease-targeted by ApoI and, in subset, by EcoRI (5’-GAATTC-3’). Unmethylated 5’-GANTC-3’ is endonuclease-targeted by HinfI and, in subset, by TfiI (5’-GAWTC-3’; W = A or T). Under current assumptions, DNA from organisms that methylate these motifs should resist against the action of the listed endonucleases.

One such clinically relevant genus is *Campylobacter* (5’-RA<u>**A**</u>TTY-3’) which is known to cause widespread foodborne illness across the globe, and is estimated to cause more than 1.5 million infections per year in the US (https://www.cdc.gov/campylobacter/faq.html) and close to nine million in the European Union (https://www.efsa.europa.eu/en/topics/topic/campylobacter). *Campylobacter* is often associated with the contamination and microbiome of poultry and wild birds (38). Theoretically, the use of ApoI and EcoRI should be able to enrich for these bacteria in metagenomic samples.

Another scenario where enrichment of pathogenic DNA for whole genome sequencing purposes is particularly useful is in the case of nosocomial infections (infections that originate in the hospital). These are often spread through patient-to-patient contact or patient-to-surface-to-patient contact and need to be traced to origin (39). Such is the case, for example, for the opportunistic pathogen *Acinetobacter baumanii* (5’-RA<u>**A**</u>TTY-3’) which initially cropped up in medical military facilities and quickly spread to civilian medical facilities by way of infected soldiers being transported through them (40). This is in addition to bacteria such as *Klebsiella pneumoniae* that may use the Dam/Dcm systems described above (41).

*Mycoplasma bovis* (5’-G<u>**A**</u>NTC-3’) is known to infect cattle and has resulted in an estimated loss of $108 million in the US annually (42). Similarly, species *abortus, melitensis* and *suis* of the genus *Brucella* (5’-G<u>**A**</u>NTC-3’) are known to cause Brucellosis in livestock. This method may accordingly prove useful in disease tracking within livestock settings.

*Oliveira and Fang et al*. detail the presence and distribution of different methylated motifs across clades of bacteria which may prove useful for a user of this method to select appropriate restriction enzymes for their organism of interest (19).

Although the patterns of modification are quite consistent in many species, we note a limitation that strains with different patterns of modification exist for some species and should be considered as potential confounders of any generalized approach. For example, B strain *E. coli* have lost their ability for Dcm methylation likely in the laboratory (43, 44). We encountered this as we tried to enrich for OP50 *E. coli* DNA and found, instead, that it was digested by PspGI and EcoRII indicating that it was not Dcm methylated. We soon learned that OP50 is derived from B strain *E. coli* (45). Also to be noted is that while Dam methyltransferases are indeed widespread (though not ubiquitous (46)) within *Gammaproteobacteria*, Dcm methyltransferases are confined to genera closely related to *Escherichia* which may serve as an advantage or disadvantage depending on the organismal DNA to be enriched for (40). Since the Dam motif is shorter than the Dcm motif, it is found more frequently in any given genome. Hence, MboI contributes most to the segregation of methylated and unmethylated DNA at these sites as compared to PspGI and EcoRII (Fig 2B) suggesting that an MboI only digestion would be sufficient to achieve strong enrichment. Indeed, it has also been found that Dam serves a core function for gene expression of virulence factors and that Dam inhibition attenuates virulence and pathogenicity in Dam bacteria in vivo (15, 18, 43). That said, pathogenic strains leading to outbreak such as O157:H7 have classically contained the genes for both Dam and Dcm (11, 47).

Of potential interest in understanding the results of REMoDE assays are the characteristics of DNA fragments from non-methylated organisms that remain after digestion and are represented in the sequencing data. Several features could result in the survival of these fragments including a lack of restriction sites in long stretches of a genome, circular DNA (that does not contain the corresponding restriction sites and is insusceptible to exonuclease degradation), or protection of DNA ends on linear fragments due to specific chemical structures or linkage to a terminal protein. Likewise novel DNA modifications (or damaged bases) could render some or all fragments from a given experimental source resistant to the initial endonuclease digestions.

Restriction-modification systems evolved such that a host cell’s restriction enzymes would be unable to digest host DNA due to the presence of protective modifications which infecting phage DNA would not have. As such, Type II restriction enzymes are very specific to their cognate restriction sites but are blocked by these modifications. This proves a useful method to distinguish modified DNA from unmodified DNA. In some cases, these enzymes are unable to cleave DNA with other modifications within the restriction site and not just with the modification associated with the corresponding restriction-modification system. Indeed, certain phage (T4, S2-L etc.) modify all instances of a base (C in T4, A in S2-L) in their genome and when purified DNA from these phages is treated with restriction enzymes, the DNA withstands the action of these enzymes (33, 34, 48). This is why a substantial enrichment of T4 DNA was observed in our experiments when there was an inadvertent inclusion of T4 DNA in our yeast DNA sample. REMoDE could therefore also be used to screen environmental samples for DNAs resistant to the action of a selection of endonucleases. Such sequences suggest the presence of non-canonical bases or modifications. This DNA could then be sequenced either by standard short read sequencing (e.g. Illumina) or by methods conducive to distinguishing modified residues such as Oxford Nanopore or PacBio Single Molecule Real Time (SMRT) sequencing.

## Methods

### Genomic DNA preparation

#### E. coli

*Protocol adapted from Green and Sambrook et al* (49). 1.5mL of an overnight culture (2x TY media) of Top10 *E. coli* was centrifuged at room temperature for 30 seconds and the supernatant removed by aspiration. 400μL of 10 mM Tris 1mM EDTA (TE) buffer at pH 8.0 was added to the tube and the bacterial pellet was resuspended via gentle vortexing. 50μL of 10% SDS and 50μL of Proteinase K (20 mg/mL in TE, pH 7.5) was added to the tube and left to incubate at 37°C for 1 hour. The digested lysate was pipetted up and down three times with a p1000 pipette to reduce viscosity. 500μL of a 1:1 mixture of phenol:chloroform (phenol equilibrated with 10mM Tris-HCl, pH 8.0) was added to the tube and pipetted up and down multiple times to mix. The mixture was then transferred to a 2mL phase lock light tube (5 PRIME 2302800) and centrifuged at 16,000 RCF at room temperature for 5 minutes. The aqueous phase was transferred to a new phase lock tube and the 1:1 phenol:chloroform extraction was repeated. The aqueous phase was then extracted twice with 500μL chloroform. The suspension was transferred to a fresh microcentrifuge tube and 25μL of 5M NaCl followed by 1mL of ice-cold 95% ethanol was added. The mixture was pipetted up and down multiple times and then centrifuged at 21,000 RCF at 4°C for 10 minutes. The supernatant was carefully removed with a pipette and left to dry for 10 minutes. The damp-dry pellet was dissolved in 100μL of TE. 2.5μL of RNaseA (10mg/mL; Thermo Scientific EN0531) was added to the solution, mixed, and left to incubate for 30 minutes at 37°C. 40μL of 5M ammonium acetate and 250μL of isopropanol were added to the mixture, mixed with a pipette and left to incubate at room temperature for 10 minutes with the cap closed. The tube was centrifuged at 21,000 RCF at room temperature for 10 minutes. The pellet was washed twice with 70% ethanol and then the ethanol was aspirated carefully with a pipette. The tube was left to dry for 10 minutes. The pellet was then dissolved in 100μL TE (pH 8.0) and left to incubate overnight at 37°C for complete dissolution. The concentration was determined using Qubit BR dsDNA reagents and a Qubit 2.0 fluorometer.

#### C. elegans

Worms from three 60mmx15mm starved plates of N2-strain (PD1074) *C. elegans* were collected by washing them off the plate with 1.5mL of 50mM NaCl and into a 1.5 mL tube. The tube was centrifuged for 40 seconds at 400 RCF at room temperature. Approximately 1200μL of the supernatant was aspirated out, leaving roughly 300μL of worms and solution. In a fresh 1.5mL tube 1.2mL of 50mM NaCl containing 5% sucrose was added. The remaining 300μL of the worms and solution was mixed and layered over the sucrose cushion. The tube was centrifuged for 40 seconds at 400 RCF. The supernatant was removed and the pellet was washed with 1.5mL of 50mM NaCl. The tube was centrifuged again for 40 seconds and the supernatant removed. 450μL of Worm Lysis Buffer (0.1M Tris pH 8.5, 0.1M NaCl, 50mM EDTA and 1% SDS) was added to the tube along with 20μL of proteinase K (20mg/mL). The tube was gently vortexed. The tube was left to incubate at 62°C for 45 minutes and vortexed four to five times throughout the incubation. 500μL of phenol was added to the tube, mixed thoroughly by pipetting up and down, and transferred to a phase lock light tube. The tube was centrifuged for 5 minutes at 16,000 RCF. The aqueous phase was transferred to a new phase lock tube and extracted with 500μLs of 1:1 phenol:chloroform. Finally, the aqueous phase, again, was extracted with 500μLs of chloroform and transferred to a fresh 1.5mL tube. 80μL of 5M ammonium acetate was added to the solution. 1mL of ethanol was added to the tube and mixed thoroughly by pipetting. The tube was then centrifuged for 5 minutes at 21,000 RCF at room temperature and the pellet was washed once with 0.5mL of ethanol and centrifuged again. The ethanol was aspirated out and the pellet was left to dry for 10 minutes after which 25μL of TE (pH 8.0) was used to resuspend it. The concentration was determined using Qubit BR dsDNA reagents and a Qubit 2.0 fluorometer. Note that RNase was not used in this preparation and thus downstream experiments with *C. elegans* contain *C. elegans* RNA, however DNA was RNaseA treated before loading onto gel in Fig 2A.

#### S. cerevisiae

4mL of an overnight S288C yeast culture (YPD media) was pelleted at 16,000 RCF for 1 minute and resuspended in 250ul of Breaking Buffer (2% (v/v) Triton X 100, 1% (w/v) SDS, 100mM NaCl, 10mM Tris base pH 8, 1mM EDTA). Approximately to the volume of 200μL of 0.5mm glass beads was added to the mixture as well as 500μL of 1:1 phenol:chloroform. The tube was vortexed, at max speed, at 4°C for 10 minutes. It was then centrifuged at 16,000 RPM for 10 minutes. 400μL of the aqueous phase was transferred to a fresh 1.5mL tube. 1μL of RNase A (10mg/mL) was added to the mixture and it was left to incubate at 37°C for 10 mins. 750 μL of 1:1 phenol:chloroform was added to the tube and mixed well with a pipette. The solution was transferred to a 2mL phase lock light tube and centrifuged for 5 minutes at 16,000 RCF. The aqueous phase was then transferred to a fresh 1.5mL tube and 65μL of 3M sodium acetate was added to the tube. 1mL of ice-cold ethanol was added to the tube, mixed, and left to incubate for 10 minutes at −20°C. The tube was centrifuged for 10 minutes at 21,000 RCF. The supernatant was carefully aspirated out and the pellet was washed with 1mL of ice-cold 70% ethanol. The tube was spun again for 10 minutes at 21,000 RCF. The supernatant was carefully aspirated out and the pellet was left to dry for 10 minutes. It was then resuspended in 20μL of TE. The concentration was determined using Qubit BR dsDNA reagents and a Qubit 2.0 fluorometer.

#### ZymoBIOMICS MCS-HMW DNA

ZymoBIOMICS MCS-HMW DNA (D6322) was obtained from Zymo Research. The concentration was determined using Qubit BR dsDNA reagents and a Qubit 2.0 fluorometer and found to be slightly lower than the manufacturer specifications. For all following experiments, the qubit-measured concentration was used instead of the manufacturer provided one.

### Endonuclease and exonuclease treatment

Note that for initial experiments, a substantial amount of DNA was used (750ng) as input and it was later found that the input could be decreased manifold. In the ZymoBIOMICS MCS-HMW DNA experiments, 75ng of input DNA was used.

Both PspGI and EcoRII target Dcm sites and were used in these experiments. The redundancy is due to both enzymes requiring conditions that were inconvenient: PspGI has a high optimal temperature which is 75C that could be detrimental to the nucleic acids in the sample and EcoRII requires the presence of two Dcm sites in close proximity for cleavage.

### *E. coli* and *C. elegans* mixture

To set up 37.5μL reactions, 1:3 mixtures of *E. coli* and *C. elegans* DNA were made by mixing 187.5ng of *E. coli* genomic DNA with 562.5ng of *C. elegans* genomic DNA in 8-strip PCR tubes. A volume of ultrapure water needed to make the reaction up to 37.5μL after the addition of rCutsmart (NEB B6004S) and PspGI (NEB R0611) was added to each reaction followed by 3.75μL of 10x rCutsmart buffer and 0.6μL (6U) of PspGI. The tubes were mixed via gentle vortexing after every step. Each mixture was incubated at 50°C for 30 minutes after which 0.6μL (3U) of MboI (NEB R0147) was added to each reaction. Each mixture was incubated at 37°C for 30 minutes after which 0.94μL of 2M NaCl was added to each reaction (to bring the NaCl concentration to roughly 50mM which is optimal for EcoRII (Thermo Scientific ER1921)). Then, 0.6μL (6U) of EcoRII was added to each reaction. The mixture was incubated at 37°C for 1 hour. The tubes were put on ice and 0.4μL (4U) of T5 exonuclease (NEB M0663) was added to each reaction and incubated for 2, 5, 10 or 20 minutes at 37°C and immediately quenched with 8μL 6x NEB purple loading dye (NEB B7024S) supplemented with 6mM EDTA (to make the total EDTA concentration in the stock tube 66mM). 12μL of the mixture was resolved on a 1% agarose gel run at 140V for 40 minutes.

74μL of ultrapure water was added to the remaining sample (to make up to 100μL total volume) and each reaction was then purified using the Zymo Genomic Clean and Concentrate kit (Zymo D4011). The DNA was eluted with 10mM Tris buffer heated to 63°C and incubated for a few minutes. The concentration was determined using Qubit HS dsDNA reagents and a Qubit 2.0 fluorometer. For control reactions, enzymes were replaced with an equal volume of ultrapure water at the appropriate point in the protocol.

### ZymoBIOMICS MCS-HMW

For the experiment in Fig 3A, 75ng of ZymoBIOMICS MCS-HMW DNA was used. A volume of ultrapure water needed to make the reaction up to 37.5μL after the addition of rCutsmart and PspGI was added to each reaction followed by 3.75μL of 10x rCutsmart buffer and 0.6μL (6U) of PspGI. The tubes were mixed via gentle vortexing after every step. Each mixture was incubated at 50°C for 30 minutes after which 0.6μL (3U) of MboI was added to each reaction. Each mixture was then incubated at 37°C for 30 minutes after which 0.94μL of 2M NaCl added to each reaction (to bring the NaCl concentration to roughly 50mM which is optimal for EcoRII). Then, 0.6μL (6U) of EcoRII was added to each reaction. The mixture was incubated at 37°C for 1 hour. Note that as shown in Fig S1, this step may be omitted and after incubation with MboI, the reaction may proceed directly to T5 exonuclease digestion. The tubes were put on ice and 0.4μL (0.4U) of T5 exonuclease diluted 1:10 in 1x NEBuffer 4 was added to each reaction and incubated for 5 minutes at 37°C and immediately quenched with 8μL 66mM EDTA. The tubes were vortexed. 52μL of TE was added to each reaction to make up to 100μL total volume and purified using the Zymo Genomic Clean and Concentrate kit. The DNA was eluted with 15 μL of 10mM Tris buffer heated to 50°C and incubated for a few minutes. This experiment was done in biological duplicate.

For the experiment in Fig 3B, the same protocol was followed as in Fig 3A except after the EcoRII incubation, 8μL of NEB Purple Loading Dye was added and the samples were loaded into a 1% agarose gel and run for 45 minutes at 120V. The high molecular weight bands were excised with a razor and dissolved in 3 volumes of Zymo Agarose Dissolving Buffer (Zymo D4001) at 50°C. They were then processed through the Zymo Genomic Clean and Concentrate kit as in Fig 3A excluding the step of the addition of ChIP DNA binding buffer. This experiment was done in biological duplicate and one of the duplicates was prepared from a previous experiment.

For the experiment in Fig 4A, the same protocol was followed as in the experiment in Fig 3A save for using either 37.5, 7.5 and 0.75ng of input ZymoBIOMICS MCS-HMW DNA.

For the experiment in Fig 4B, different ratios of ZymoBIOMICS MCS-HMW DNA and *S. cerevisiae* DNA were mixed together 1:1 (37.5ng:37.5ng), 1:9 (7.5ng:67.5ng) and 1:99 (0.75ng:74.25ng) and the same protocol was followed as in Fig 3A save for the T5 exonuclease incubation being 20 minutes instead of 5. This experiment was done in biological duplicate.

For the experiment in Fig 5, the same protocol was followed as in Fig 3A, save for a 1 hour incubation with 0.3μL (3U) of DpnI (NEB R0176) at 37°C instead of using PspGI, MboI or EcoRII.

## Library Preparation and Miseq Sequencing

Nextera XT (Illumina FC-131-1024) library preparation was used to build sequencing libraries for all experiments. One-third of the recommended volumes of the manufacturer protocol were used i.e. 3.33μL of Tagmentation buffer, 1.67μL of 0.2ng/μL DNA, 1.67μL of Tn5 mix, 1.67μL of the neutralizing buffer, 1.67μL of each index followed by 5μL of the polymerase mix. The transposition incubation was done at 37°C for 5 minutes. Amplification was performed as in the Nextera XT protocol with 12 cycles of amplification. The amplified libraries were resolved on a gel and DNA of the range of 300 to 600 bp was excised for gel recovery using the Zymo gel extraction kit (Zymo D4007). Concentrations of DNA were determined with Qubit HS dsDNA reagents and a Qubit 2.0 fluorometer. The libraries were pooled and sequenced on an Illumina Miseq sequencer; 78 cycle, paired-end.

## Data Analysis

Reads were demultiplexed and mapped to their corresponding genomes via Bowtie2 (50) using default settings. The reference file in each experiment contained the genomes of each organism whose DNA was used in the experiment. Mapped reads for each organism were counted and plotted using matplotlib in Python3 on Jupyter Notebook. Relative enrichment was calculated as follows:

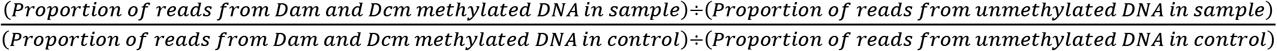

### Genome sequences

For *E. coli* and *C. elegans* experiments, genome assemblies GCA_000005845.2 (GenBank) and UNSB01000000 (European Nucleotide Archive) were used respectively. These genomes were combined into a single FASTA file used as a reference for Bowtie2 and the alignments were output as SAM files.

For ZymoBIOMICS MCS-HMW DNA experiments, genome assemblies were obtained from the protocol of this reagent (https://s3.amazonaws.com/zymo-files/BioPool/D6322.refseq.zip). The genomes were combined into a single file. Since the assembly that was included for *S. cerevisiae* was heavily discontiguous and since the S288C strain of *S. cerevisiae* was used in the experiment for Fig 4B, the provided *S. cerevisiae* was instead replaced with the latest assembly available on the Saccharomyces Genome Database (S288C_reference_sequence_R64-3-1_20210421). Also added to this file was the genome for T4 phage (OL964735.1; GenBank) as this sequence appears in some sequencing datasets due to use in other ongoing experiments. Finally, for some samples, reads that did not map to any of the listed genomes were assembled using SPADES with default parameters. When these contigs were input into Blastn it revealed the presence of the aforementioned T4 phage DNA but also plasmids of *S. cerevisiae*, and *S. enterica* that were not included in the reference genomes (*S. cerevisiae:* CP059538.1, J01347.1; *S. enterica:*CP012345.2; GenBank). These plasmids were also added to the reference genome file.

### *In silico* digest of *C. elegans* genome

The *C. elegans* genome was digested *in silico* based on sites where a Dam or Dcm cleavage site is expected. Each read mapped by Bowtie2 was located to the theoretical fragment by genomic coordinates. The theoretical length of the containing fragment(s) for each read was assessed by measuring the number of bases between upstream and downstream cut sites.

## Acknowledgements

We thank the lab of Dr. Gavin Sherlock (Stanford University) and A. Pyke for providing the S288C strain of *S. cerevisiae*. We also thank K. Artiles, M. McCoy, O. Ilbay, L. Wahba, M. Shoura, D. Jeong, E. Greenwald, D. Galls, A. Straight, P. Sidhwani, K. Sundararajan, C. Limouse, K. Fryer, R. Brown, O. Smith, R. Ladurner and M. Gebala for discussion on the project. Support for this work was provided by NIH Grant R35GM130366. SUE was supported by NIH Grant T32HG000044. JLC was supported by the intramural research program of the National Library of Medicine, National Institutes of Health. The opinions expressed in this article are those of the author and do not reflect the view of the National Institutes of Health, the U.S. Food and Drug Administration, the Department of Health and Human Services, or the United States government. Figures were created with BioRender.com.

